# Disentangling the contribution of each descriptive characteristic of every single mutation to its functional effects

**DOI:** 10.1101/867812

**Authors:** C. K. Sruthi, Meher K. Prakash

**Affiliations:** Theoretical Sciences Unit, Jawaharlal Nehru Centre for Advanced Scientific Research, Bangalore 560064, India

**Keywords:** Mutational effects, Deep mutational scan, Protein design, Conservation, Artificial Intelligence, Interpretable AI

## Abstract

Mutational effects predictions continue to improve in accuracy as advanced artificial intelligence (AI) algorithms are trained on exhaustive experimental data. The next natural questions to ask are if it is now possible to gain insights into which attribute of the mutation contributes how much to the mutational effects, and if one can develop universal rules for mapping the descriptors to mutational effects. In this work, we mainly address the former aspect using a framework of interpretable AI. Relations between the physico-chemical descriptors and their contributions to the mutational effects are extracted by analyzing the data on 29,832 variants from 8 systematic deep-mutational scan studies. It is found that the intuitive dependences of fitness and solubility on the distance of the amino acid from active site could be extracted and quantified. The dependence of the mutational effect contributions on the number of contacts an amino acid has or the BLOSUM score descriptor of the change showed universal trends. Our attempts in the present work to explain the quantitative differences in the dependence on conservation and SASA across proteins were not successful. The work nevertheless brings transparency into the predictions, development of rules, and will hopefully lead to uncovering the universalities among these rules.

## Introduction

Proteins perform the most critical cellular functions, and yet they are very delicate in their design. A small perturbation to the sequence such as a change in a single amino acid of the protein, known as a mutation, can alter their structure and function significantly. In fact, several of the disease phenotypes such as cancers, Alzheimer’s^1^ as well as other problems associated with infectious diseases such as antibiotic resistance^2^ can all be traced back to mutations in proteins. Understanding the effects of mutations is also relevant in protein engineering, where one would like to introduce the changes that affect the solubility and function of the proteins in a predictable way.

One of the ways of understanding mutational effects is by introducing these mutations in the proteins,^3^ in a site-directed or random fashion, and studying the structural or functional consequences of it. In a commonly practised protocol, known as the alanine scanning,^4^ amino acids of interest are substituted by alanine, which is small and neutral, and the functional consequences are studied. In recent years, technological advances allowed a much more detailed scan known as the deep mutational scanning (DMS). ^5–7^ In DMS, each amino acid is replaced by all possible 19 amino acids, and cellular level consequences are studied. The exhaustive sampling allows thousands of mutations to be studied at a single substitution level, and hundreds of thousands with two to three simultaneous substitutions.

The computational efforts to understand the mutations are also making significant strides. Artificial Intelligence based models are useful for complementing the missing data in DMS, ^8,9^ or making detailed predictions of the complete mutational scanning using co-evolutionary relations^10,11^ or by training on a few DMS experiments.^12^ The models trained on large mutational databases curated from across the literature^13^ or from a few DMS studies^12^ are making good predictions of the mutational effects. The wealth of data that is being generated through the DMS experiments are also useful for benchmarking or evaluating different predictors.^14,15^

Ability to make reliable mutational effects predictions using machine learning approaches brings one to a natural point of asking the questions that are being asked in other areas where machine learning is used, such as to why the predictions should be trusted^16^ or alternatively if the predictions can be interpreted. ^17,18^ The philosophical debate about predictability versus interpretability^19^ is being revisited in several areas of machine learning, specifically when one is interested in controlling the effects by developing an understanding for the contributing factors. We introduce this notion of interpretability or explainability in the context of mutational effects prediction. Interpretability begins with a small shift in perspective, from asking how do various factors contribute to the *set* of predictions, to what is the contribution of the various factors to an *individual* prediction. In a linear regression model, knowing the measured outcome, it is trivial to understand the relative contributions from the different factors. The same is not true while working with machine learning models, which have non-trivial and non-explicit relations between the inputs and the outcome.

We use an approach known as the SHAP (SHapley Additive exPlanations)^20,21^ that is being used in several areas of machine learning, to interpret the contributions to the mutational effects. SHAP is based on the game theoretical questions raised by Shapley ^22^ about how the gain can be shared by different contributing players. In SHAP, the feature contributions are additive, thus making their relation to the outcome easy to interpret. We apply SHAP to interpret the outcomes of fitness^23^ and solubility^24^ in the deep mutational scanning studies of β-lactamase protein. The interpretability allows us to revisit the classical intuitions on how different factors can influence mutations from a quantitative perspective, and begin a search for universal rules for understanding mutational effects.

## Methods

### Deep mutational scanning data

The present work analyzes 29,832 mutational effects from 8 different deep mutational scanning studies (Supplementary Table 1). The mutational effects of β-lactamase on fitness was obtained from Stiffler et al^23^ and solubility from Kle-smith et al.^24^ We use the yeast surface display (YSD) data on the effects of mutations on solubility^24^ and the changes in relative fitness of *E. coli* when a mutant containing strain is challenged with 2500 *µg/ml* ampicillin.^23^ The analyses on the solubility were presented for the substitutions at positions 61-215,^24^ with protein data bank identity 1M40,^25^ and for consistency, we chose to work with the same set of mutations both for the solubility and cellular fitness. For APH(3^‘^)-II, Hsp90 and MAPK1 the mutational effect scores available as relative fitness in the study of Gray et al.^26^ were used.

The fitness scores (*F*_*i*_) obtained from the different studies were rescaled by the overall effect sizes in the assay, to allow a direct comparison across proteins. The rescaling was performed as: 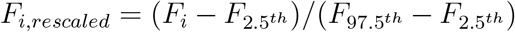, where 2.5^*th*^ and 97.5^*th*^ denote the 2.5 and 97.5 percentiles respectively of the fitness score within that data set. This will result in scores that roughly range from 0 to 1 and an increase in the score corresponds to an increase in fitness. Unlike for other proteins, the measurements on MAPK1 had an opposite meaning with positive values of the relative fitness referring to deleterious mutations. For consistency of notation among the different proteins, the fitness scores were multiplied by (−1) before rescaling.

### Descriptive features for AI model

We used 12 physico-chemical descriptive features to describe the nature of the wild-type amino acid as well as the substitution:^9^

#### Structural factors

1. *SASA* - Solvent accessible surface area was calculated with the function *gmx sasa* of GROMACS with default parameters 2. *SS* - Based on the secondary structural element the wild-type amino acid belongs to, a score of 1 was given if it is ordered (helix or sheet) and 0 otherwise. 3. *Contacts* - number of neighboring amino acids within 4 Å of the wild type amino acid, 4. *Catalytic_Dist*, which is the distance of the amino acid from the closest catalytic site. This feature was used only in the analysis of β-lactamase for comparisons with the solubility studies,^24^ 5. *Avg_Commutetime*, which is the average commute time reflecting how well the amino acid is connected to the rest of the protein and was calculated following the article of Chennubhotla et al.^27^ The protein data bank (PDB) identities for the structures used in the case of each protein are given in Supplementary Table 1.

#### Sequence factors

6. *BLOSUM*, BLOSUM65 substitution matrix score,^28^ 7. *HB_WT*, hydrophobicity of the wild-type amino acid according to Kyte-Dolittle scale,^29^ 8. *HB_MUT*, hydrophobicity of the amino acid after mutation.

#### Multiple sequence alignment derived factors

The multiple sequence alignments from Pfam^30^ were used for the proteins β-lactamase, Bgl3 and LGK, the Pfam IDs being PF13354, PF00232 and PF03702 respectively. For the other proteins the sequences obtained through PSI-BLAST search were aligned using Clustal Omega.^31^ The alignment was truncated to the sequence of reference protein. All sequences with more than 20% gaps were removed from the alignment. This alignment was used to calculate:

9. *PSSM_WT* representing the position specific scoring matrix (PSSM) score of the wild-type amino acid using PSI-BLAST with default parameters, 10. *PSSM_MUT* - PSSM score of the mutant, 11. *Conservation* - frequency of occurrence of the most frequent residue at a position.

#### Co-evolutionary factors

12. *Avg_Corr* for a given position the feature was calculated as the average of co-evolutionary correlation of an amino acid with all other amino acids as follows. From MSA, the correlated evolution of amino acids was quantified using the statistical coupling analysis following Halabi et al.^32^ For that the alignment was binarized by replacing the most frequent amino acid at a position with 1 and all others with 0. The co-evolution between amino acids *i* and *j* was then calculated using the equation: *C*_*ij*_ = *ϕ*_*i*_*ϕ*_*j*_ | ⟨*x*_*i*_*x*_*j*_⟩_*s*_ − ⟨*x*_*i*_⟩_*s*_ ⟨*x*_*j*_⟩_*s*_|. *ϕ*_*i*_ is defined as 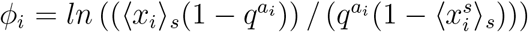, where 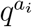 is the probability of occurrence of the conserved residue *a*_*i*_ at position *i* among all proteins. *x*_*i*_ is the *i*^*th*^ residue position in the binarized MSA and ⟨ ⟩_*s*_ denotes the average over sequences.

### AI model

The descriptive features of the contextual nature of each mutation in a given protein were related to the to the rescaled fitness (*F*_*i,rescaled*_) using AI models. The training was independently performed for each protein data set. The analyses were performed in Python 3.7, using the *XGRegressor* implementation of the *XGBoost* ^33^ package. 75% of the mutational data was used for training and the remaining 25% for predictions. For each data set, the learning rate was fixed at 0.01 and first the hyper parameter *n_estimators* was optimized using the *cv* function of XGBoost with the early stopping algorithm. The parameters *max_depth, min_child_weight, gamma, subsample, colsample_bytree* and *reg_alpha* were then tuned using the GridSearchCV function of scikit-learn in multiple steps. In the first step optimal values for *max_depth* and *min_child_weight* were obtained following which *gamma* was tuned. Next step involved optimization of the parameters *subsample* and *col-sample_bytree*. Finally models were developed with different *reg_alpha* values to find the find the parameter value that improves model’s predictive power. At each step of this optimization all other parameters except the ones that are being tuned were kept constant. Once the optimal value of a parameter is determined, that value was used in all the subsequent steps. The steps of this optimization and the grids defined for each parameter are summarized in Supplementary Table 2. The parameters that yielded the least average Mean Squared Error (MSE) for the test sets of 5-fold cross-validation analysis were chosen. The optimal parameters obtained are given in Supplementary Table 3. The models trained with these optimized parameters were further analyzed using SHAP. For all proteins a 5-fold cross-validation analysis was performed to verify the robustness of the SHAP values obtained as the training set is changed.

### Interpretable AI model

SHapley Additive explanation (SHAP) uses the formalism where an explanation model *g* is defined in terms of the parameter set *z*_*i*_’ defining each instance (in our case each individual mutation) and their corresponding additive contribution *ϕ*_*i*_ weights.^20,21^

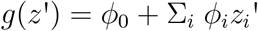

The explanatory model is subject to three conditions known as:

*Local accuracy* - which ensures that it matches the calculated effect *f* (*z*) when *z*’ = *z*, i.e., *g*(*z*’) = *f* (*z*) when *z*’ = *z*,

*Missingness* - which ensures that if a variable is *z*_*i*_’ = 0, then the weight corresponding to it is *ϕ*_*i*_ = 0.

*Consistency* - when an input’s contribution increases or stays the same regardless of the other inputs, then its weight should not decrease.

By solving for these three conditions, one obtains the SHAP contribution weights corresponding to each individual input instance. We used the SHAP implementation for performing the interpretable AI calculations (https://github.com/slundberg/shap),where contributions of each factor to every mutational effect calculation are determined. The results presented in this work discuss these SHAP weight factors.

## Results and Discussion

### Decoupling the contributions of each factor to individual mutational effects

The focus of this work was to go one step beyond accurate predictions of the relative fitness changes due to mutations, by decoupling the contributions of individual factors to each single mutational effect prediction. The deep mutational scanning data from eight proteins were analyzed: β-lactamase,^23^ APH(3^‘^)-II,^34^ Hsp90,^35^ MAPK1,^36^ UBE2I,^8^ TPK1,^8^ Bgl3^37^ and LGK.^38^ In addition, an additional data set on the deep mutational scan of solubility changes due to mutations in β-lacatmase was also analyzed. Separate AI models were developed for each of these protein data sets, to relate the physico-chemical descriptive variables from structural and sequence origins to the mutational outcomes (**Methods**). Training on 75% of the data from each protein resulted in good quality of predictions. For example, the predictions of the fitness and solubility effects had Pearson correlation coefficients of 0.91 and 0.79 respectively (Supplementary Figure 1) The quality of results for analyses is shown in Supplementary Table 3. Taking confidence in these predictions, we performed the decoupling analysis to obtain the contributing factors in each individual mutation using SHAP method.

Figure 1 illustrates the predictions for a specific mutation L207N and the relative contributions of the different factors. The predicted rescaled fitness (0.13) and solubility (0.25) for this mutation compare well with the experimental observations (0.13, 0.21 respectively). The interpretable aspect of the prediction is shown in the decomposition of the various factors, firstly segregated by positive and negative contributions: factors labeled in red aiding a better fitness or solubility, and those in blue having the opposite effect. The illustration shows that the hydrophobicity index of the wild-type amino acid contributes in opposite ways to the fitness and solubility. The overall effect all individual factors may also be summarized to obtain a comprehensive perspective on the contributions of the different factors to the fitness (Figure 2A) and solubility (Figure 2B). Observing the range of the values assumed, one can infer that SASA, BLOSUM and Avg_Corr contribute significantly to both solubility and fitness.

**Figure 1:**
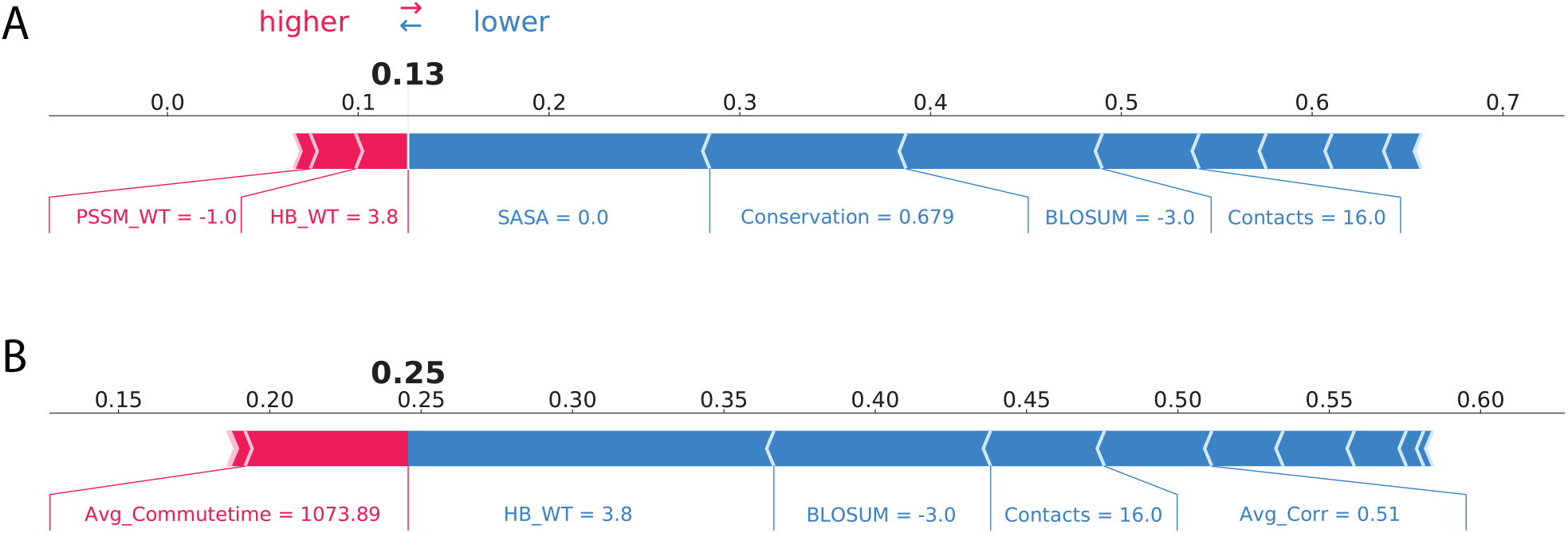
Decoupling the contributions. Illustration of the contributions of various factors to the effects of mutating Leucine in position 207 to Asparagine (L207N) in β-lactamase: A. Fitness and B. Solubility predictions. As indicated by the direction of the arrows, the factors in red contribute to an increase in the fitness or solubility and those in blue have the opposite effect. Whether a specific factor tends to increase or decrease the mutational effect depends on the individual case, as seen for example from the direction and magnitude of the contribution of the hydrophobicity of the wild-type amino acid (HB_WT). The features are labeled along with the values they assume in this specific instance, for the specific mutation. The illustrations are generated using the Python implementation of SHAP ((https://github.com/slundberg/shap)).

**Figure 2:**
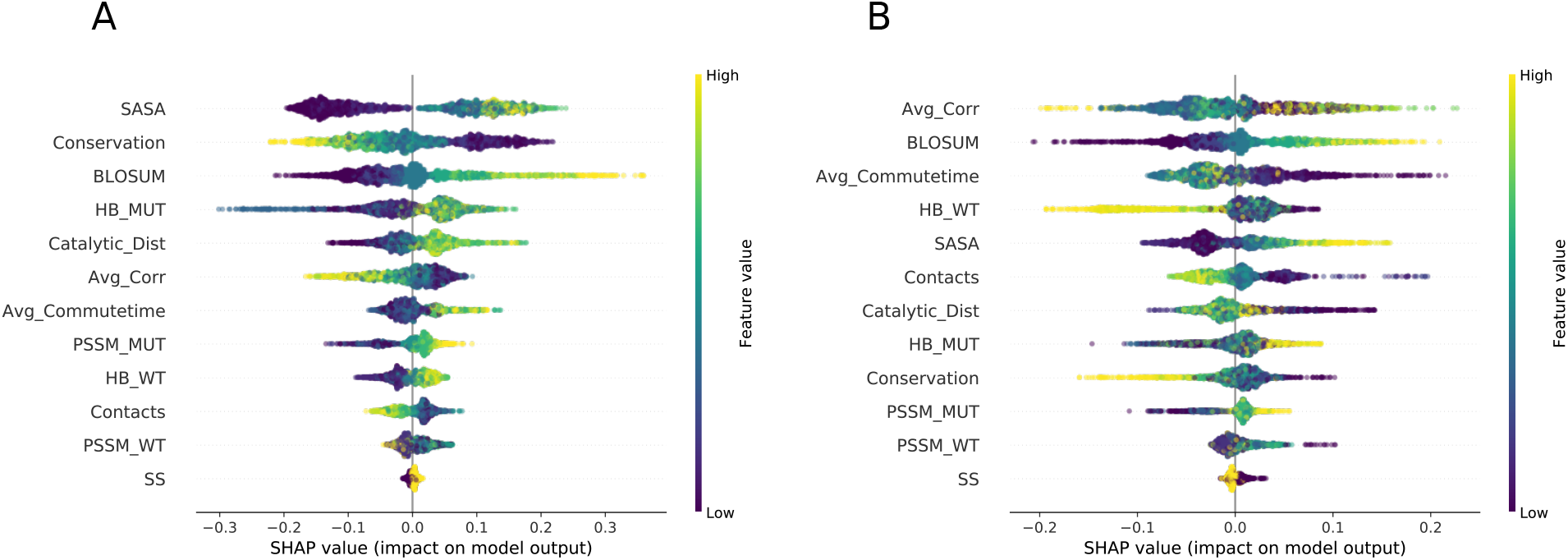
Summarizing the contributions. We analyzed the deep mutational scanning data where the consequences of any of the 19 possible amino acid substitutions at each of the positions (61-215) were measured. The individual contributions to (A.) fitness and (B.) solubility obtained from each of these mutations are summarized in the plot. Along each line, one finds the name of the feature, a distribution of the SHAP values across the complete set of mutations, along with the color indicator of the fitness and solubility outcome associated with mutation. A significant fraction of the contribution to overall mutational effect is due to the variables that describe the context of the amino acids in the wild-type, rather the variables which describe the precise nature of the mutation (Supplementary Figure 2).

### Validating decoupled contributions with experimental findings

Of all the factors that can determine the consequences of a mutation, the role of at least conservation and distance from the active site can be believed to be intuitive. However, in the naive scatter plots in Figures 3A, 3B where the outcomes are plotted against single variables, one does not see any correlation. In this projected representation of the multifactorial effect to single variable, the relation of fitness or solubility to conservation is not easy to infer. Klesmith et al.^24^ performed a naive Bayesian classification to clarify the chance that a mutation characterized by a feature is statistically likely to be deleterious or neutral. An interesting design trade-off was observed,^24^ as the distance from the catalytic site had opposite effects on solubility and fitness. We ask if one can go beyond the classification models to quantify these dependencies using SHAP analysis.

**Figure 3:**
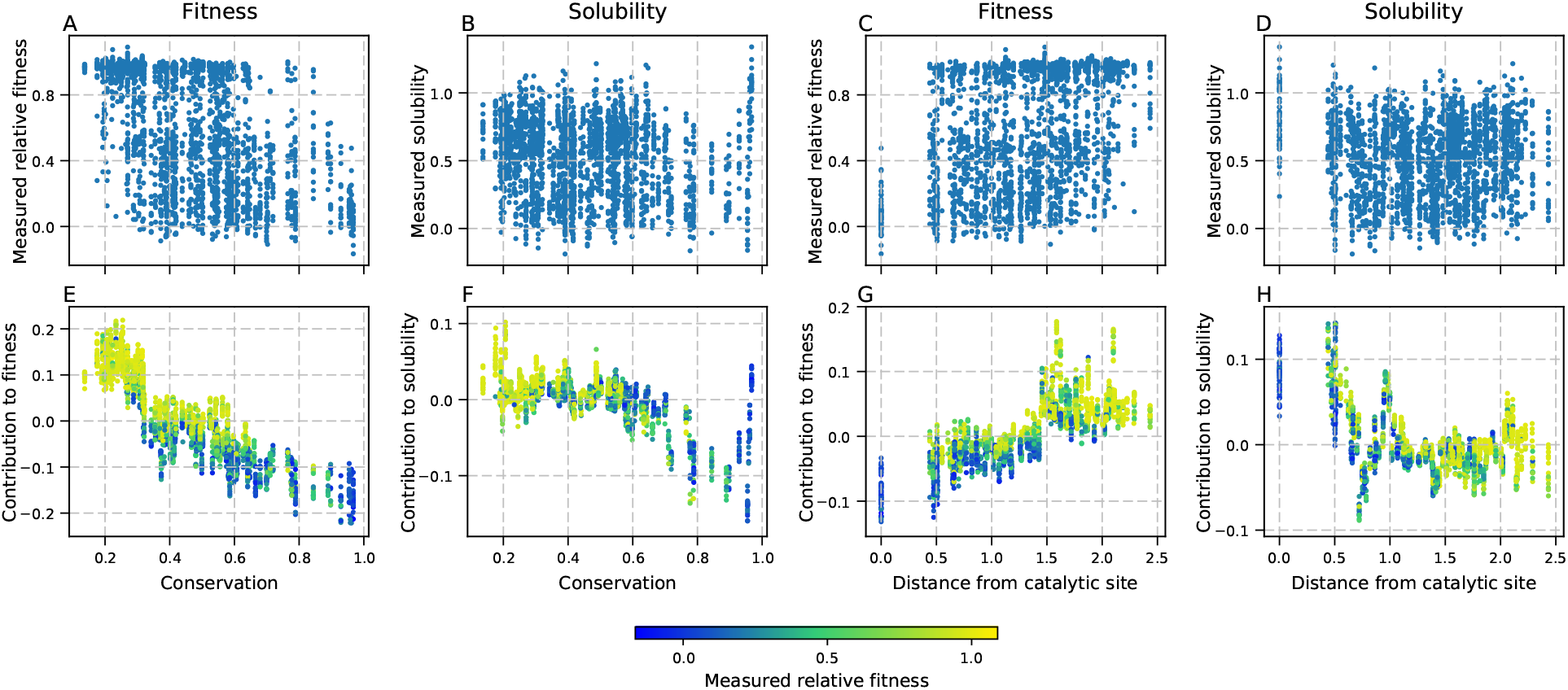
Extracting relations. The scatter plots of A-D of fitness and solubility relative to conservation and the distance from the catalytic site do not show a clear pattern of what one can expect from substituting an amino acid with high conservation or further away from catalytic site. On the contrary, E,F. the SHAP contributions highlight a very clear pattern of reducing SHAP values with increasing conservation which suggests that the fitness and solubility decrease with the substitution of a conserved amino acid. G, H. SHAP contributions show a contrasting behavior where the distance from the catalytic site has opposite effects on fitness and solubility, in line with the classifications.^24^ The colorbar represents the observed fitness changes.

The contributions from the conservation to the fitness and solubility changes of β-lactamase obtained using the SHAP analysis are plotted in Figures 3E, 3F. The intuitive patterns that are expected become apparent when the contributions from the single factor are decoupled, rather than projected. Figures 3C, 3D, 3G, 3H illustrates how the distance from the catalytic site makes predictable contributions to the fitness, and solubility. Interestingly the two outcomes (Figures 3G, 3H) show an opposite dependence on the distance from the catalytic site, as was seen using a classification model.^24^ The lower panel of Figure 3 shows a detailed quantitative relation between the contributing factors and the measurable outcomes. These comparisons with the intuitions and observations thus serve to validate the meaning of the decoupled contributions extracted using the SHAP analysis.

### Comparing the variable dependencies from multiple proteins

The deep mutational scan data predictions for the 8 proteins was analyzed using SHAP methodology. The contribution to the fitness changes when an amino acid with a known degree of conservation is mutated shown in Figure 4. Similar analyses were performed for the features Contacts, BLOSUM and SASA (Supplementary Figures 3 to 5), and the trends appeared intuitive.

**Figure 4:**
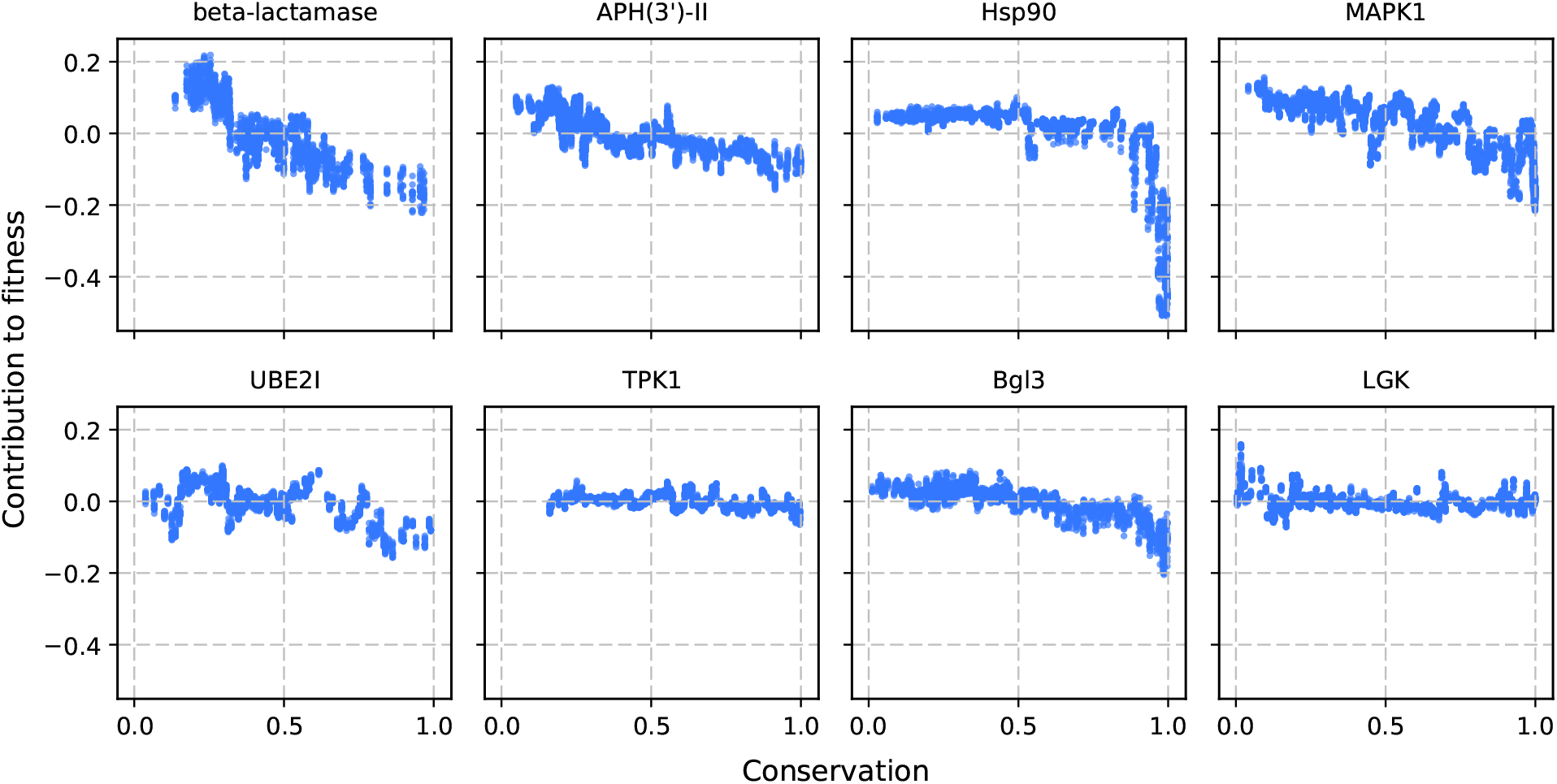
Role of conservation. The contribution to the fitness changes when an amino acid with a certain degree of conservation is substituted is shown. Results from the 8 proteins are shown.

In search of universal principles that may be valid across all proteins, we make a direct comparison in Figure 5 across proteins by overlaying the contributions to the fitness scores from the different proteins. The number of contacts an amino acid has in the wild-type protein and the BLOSUM descriptor of the change of amino acids were seen to have universal trends across the 8 proteins we studied, with similar qualitative and quantitative trends. A similar characer was not observed in relation to conservation or SASA and we tried to understand if the differences that appear in this search for universality of contributions may be addressed using one or many systemic descriptors of the proteins such as its size, polarity, degree of structural order (Supplementary Table 4). Hsp90 which showed the highest change in fitness upon mutation of highly conserved amino acids had the lowest aggregate percentage of helical and β-sheet structures. β-lactamase and Bgl3 which showed significant changes for the buried residues with SASA < 0.2 nm^2^ were also probed to see if any insights in terms of the number of contacts or the structural character of these buried residues is different for these proteins. At this stage, no correlations could be established between the observed deviations from universality and individual systemic descriptors of the proteins.

**Figure 5:**
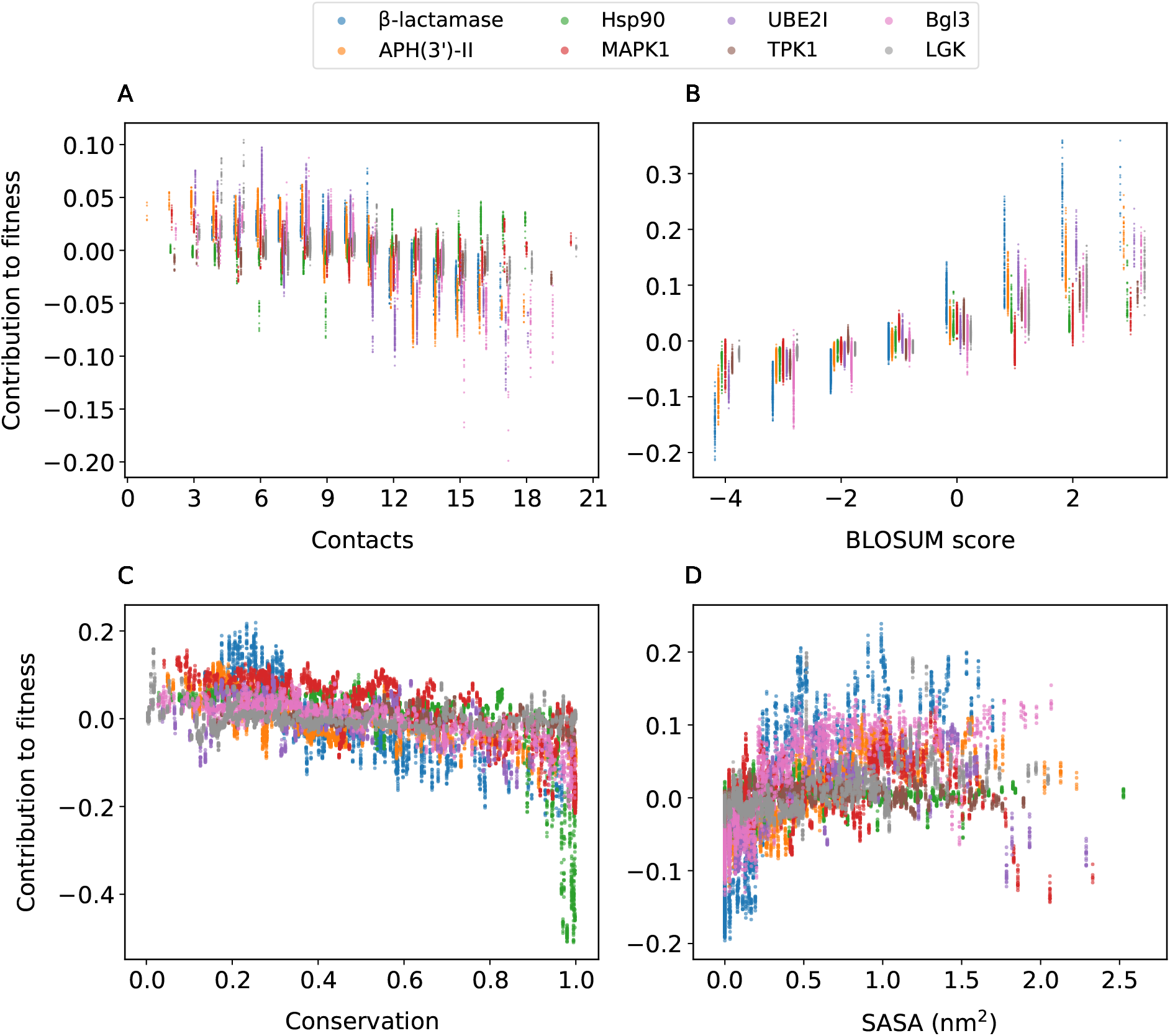
Search for universality. The contributions to fitness from the different descriptors are shown. In each of the panels, the fitness contributions from all eight proteins are overlaid. The contributions from (A) number of contacts and (B) BLOSUM substitution matrix are universal across all eight proteins. (C) Conservation and (D) SASA contributions show differences among the different proteins, and the patterns among these differences need further exploration.

### Towards simple rules for understanding mutational effects

Understanding how mutations affect proteins is thus not easy. At times a single mutation can lead to deleterious effects, and yet evolutionarily one sees homologous proteins with as little as 50% sequence identity performing similar functions. Systematic deep mutational scan experiments, accurate predictions, interpreting the contributions of different descriptors will help navigate towards this ultimate goal of developing a better understanding and designing. Using systematic mutational data from deep mutational scans, and advanced AI methods for interpretability or explainability of the predictions, we disentangled the contributions from the various descriptive factors to the overall mutational effects in this work. The process uncovered patterns relative to individual determinants which are otherwise buried in the projected representations. The approach we used is linear in nature, thus the contributions from the individual descriptive factors can be added to obtain the overall mutational effect.

This decoupling of the effects into additive contributions allowed us to make comparisons, and find universal patterns relative to BLOSUM and the number of contacts. The next goal in this approach for uncovering the rules for understanding mutational effects is to perform a meta-analysis to relate the descriptive characteristics of the different slopes or sharp drop characteristics as observed relative to conservation and SASA to overall descriptors of the protein, such as their size, structural order, etc. A meta-analysis performed over a larger set of proteins may be helpful in developing a rule-based mutational effect interpretation framework, which may subsequently bypass the AI models that led to it.

## Conclusions

The dichotomy one faces in mutational effects predictions is that on the one hand, accurate predictions are possible using AI, and on the other hand it is still hard to easily observe intuitive relations from the data that as the degree of conservation of the wild-type amino acid decreases, its contribution to the fitness continuously decreases. To address this dichotomy, we use a framework of AI to decouple the contributions from different descriptive factors. A potential outcome of this disentangling is that one may find universal dependences of the contributions from the descriptive factors across different proteins or find patterns which can be modelled using an additional meta-analysis. Both these will play an important role in understanding how mutational effects influence protein function in natural or engineering situations.

## Conflicts of Interest

The authors declare no conflicts of interest.

### Supporting Information Available

- **Supplementary Information**.**pdf:** This file contains all the supplementary figures and tables mentioned in the main text of the manuscript.
- **Scripts:** The scripts and data used in the analysis are available at https://github.com/meherpr/InterpretMutations

## References

(1) Crow, J. F. The origins, patterns and implications of human spontaneous mutation. Nature Reviews Genetics 2000, 1, 40–47.

(2) Stewart, P. S., Costerton, J. W. Antibiotic resistance of bacteria in biofilms. The lancet 2001, 358, 135–138.

(3) Fersht, A. R. Dissection of the structure and activity of the tyrosyl-tRNA synthetase by site-directed mutagenesis. Biochemistry 1987, 26, 8031–8037.

(4) Cunningham, B.; Wells, J. High-resolution epitope mapping of high-receptor interactions by alanine-scanning mutagenesis. Science 1989, 244, 1081–1085.

(5) Fowler, D. M.; Araya, C. L.; Fleishman, S. J.; Kellogg, E. H.; Stephany, J. J.; Baker, D.; Fields, S. High-resolution mapping of protein sequence-function relationships. Nature Methods 2010, 7, 741.

(6) Fowler, D. M.; Fields, S. Deep mutational scanning: a new style of protein science. Nature Methods 2014, 11, 801–807.

(7) Starita, L. M.; Young, D. L.; Islam, M.; Kitzman, J. O.; Gullingsrud, J.; Hause, R. J.; Fowler, D. M.; Parvin, J. D.; Shendure, J.; Fields, S. Massively Parallel Functional Analysis of BRCA1 RING Domain Variants. Genetics 2015, 200, 413+.

(8) Weile, J.; Sun, S.; Cote, A. G.; Knapp, J.; Verby, M.; Mellor, J. C.; Wu, Y.; Pons, C.; Wong, C.; van Lieshout, N. et al. A framework for exhaustively mapping functional missense variants. Molecular Systems Biology 2017, 13.

(9) Sruthi, C.; Prakash, M. Deep2Full: Evaluating strategies for selecting the minimal mutational experiments for optimal computational predictions of deep mutational scan outcomes. PloS one 2020, 15.

(10) Hopf, T. A.; Ingraham, J. B.; Poelwijk, F. J.; Scharfe, C. P. I.; Springer, M.; Sander, C.; Marks, D. S. Mutation effects predicted from sequence co-variation. Nature Biotechnology 2017, 35, 128–135.

(11) Riesselman, A. J.; Ingraham, J. B.; Marks, D. S. Deep generative models of genetic variation capture the effects of mutations. Nature Methods 2018, 15, 816+.

(12) Gray, V. E.; Hause, R. J.; Luebeck, J.; Shendure, J.; Fowler, D. M. Quantitative missense variant effect prediction using large-scale mutagenesis data. Cell systems 2018, 6, 116–124.

(13) Hecht, M.; Bromberg, Y.; Rost, B. Better prediction of functional effects for sequence variants. BMC genomics 2015, 16, S1.

(14) Reeb, J.; Wirth, T.; Rost, B. Variant effect predictions capture some aspects of deep mutational scanning experiments. BMC bioinformatics 2020, 21, 1–12.

(15) Livesey, B. J.; Marsh, J. A. Using deep mutational scanning to benchmark variant effect predictors and identify disease mutations. Molecular Systems Biology 2020, 16, e9380.

(16) Ribeiro, M. T.; Singh, S.; Guestrin, C. “ Why should i trust you?” Explaining the predictions of any classifier. Proceedings of the 22nd ACM SIGKDD international conference on knowledge discovery and data mining. 2016; pp 1135–1144.

(17) Datta, A.; Sen, S.; Zick, Y. Algorithmic transparency via quantitative input influence: Theory and experiments with learning systems. 2016 IEEE symposium on security and privacy (SP). 2016; pp 598–617.

(18) Štrumbelj, E.; Kononenko, I. Explaining prediction models and individual predictions with feature contributions. Knowledge and information systems 2014, 41, 647–665.

(19) Breiman, L. Statistical modeling: The two cultures (with comments and a rejoinder by the author). Statistical science 2001, 16, 199–231.

(20) Lundberg, S. M.; Lee, S.-I. A unified approach to interpreting model predictions. Advances in neural information processing systems. 2017; pp 4765–4774.

(21) Lundberg, S. M.; Erion, G. G.; Lee, S.-I. Consistent individualized feature attribution for tree ensembles. arXiv preprint 1802.03888 2018,

(22) Shapley, L. S. A Value for n-person Games, volume II of Contributions to the Theory of Games. 1953.

(23) Stiffler, M. A.; Hekstra, D. R.; Ranganathan, R. Evolvability as a Function of Purifying Selection in TEM-1 beta-Lactamase. Cell 2015, 160, 882–892.

(24) Klesmith, J. R.; Bacik, J.-P.; Wrenbeck, E. E.; Michalczyk, R.; Whitehead, T. A. Trade-offs between enzyme fitness and solubility illuminated by deep mutational scanning. Proceedings of the National Academy of Sciences 2017, 114, 2265–2270.

(25) Minasov, G.; Wang, X.; Shoichet, B. K. An ultrahigh resolution structure of TEM-1 β-lactamase suggests a role for Glu166 as the general base in acylation. Journal of the American Chemical Society 2002, 124, 5333–5340.

(26) Gray, V. E.; Hause, R. J.; Fowler, D. M. Analysis of large-scale mutagenesis data to assess the impact of single amino acid substitutions. Genetics 2017, 207, 53–61.

(27) Chennubhotla, C.; Bahar, I. Signal propagation in proteins and relation to equilibrium fluctuations. PLOS Computational Biology 2007, 3, 1716–1726.

(28) Henikoff, S.; Henikoff, J. Amino-acid substitution matrices from protein blocks. Proceedings of the National Academy of Sciences, USA 1992, 89, 10915–10919.

(29) Kyte, J.; Doolittle, R. F. A simple method for displaying the hydropathic character of a protein. Journal of molecular biology 1982, 157, 105–132.

(30) El-Gebali, S.; Mistry, J.; Bateman, A.; Eddy, S. R.; Luciani, A.; Potter, S. C.; Qureshi, M.; Richardson, L. J.; Salazar, G. A.; Smart, A. et al. The Pfam protein families database in 2019. Nucleic acids research 2019, 47, D427–D432.

(31) Sievers, F.; Wilm, A.; Dineen, D.; Gibson, T. J.; Karplus, K.; Li, W.; Lopez, R.; McWilliam, H.; Remmert, M.; Söding, J. et al. Fast, scalable generation of high-quality protein multiple sequence alignments using Clustal Omega. Molecular systems biology 2011, 7, 539.

(32) Halabi, N.; Rivoire, O.; Leibler, S.; Ranganathan, R. Protein sectors: evolutionary units of three-dimensional structure. Cell 2009, 138, 774–786.

(33) Chen, T.; Guestrin, C. Xgboost: A scalable tree boosting system. Proceedings of the 22nd acm sigkdd international conference on knowledge discovery and data mining. 2016; pp 785–794.

(34) Melnikov, A.; Rogov, P.; Wang, L.; Gnirke, A.; Mikkelsen, T. S. Comprehensive mutational scanning of a kinase in vivo reveals substrate-dependent fitness landscapes. Nucleic Acids Research 2014, 42.

(35) Mishra, P.; Flynn, J. M.; Starr, T. N.; Bolon, D. N. A. Systematic Mutant Analyses Elucidate General and Client-Specific Aspects of Hsp90 Function. Cell Reports 2016, 15, 588–598.

(36) Brenan, L.; Andreev, A.; Cohen, O.; Pantel, S.; Kamburov, A.; Cacchiarelli, D.; Persky, N. S., Zhu, C.; Bagul, M.; Goetz, E. M. et al. Phenotypic Characterization of a Comprehensive Set of MAPK1/ERK2 Missense Mutants. Cell Reports 2016, 17, 1171–1183.

(37) Romero, P. A.; Tran, T. M.; Abate, A. R. Dissecting enzyme function with microfluidic-based deep mutational scanning. Proceedings of the National Academy of Sciences 2015, 112, 7159–7164.

(38) Klesmith, J. R.; Bacik, J.-P.; Michalczyk, R.; Whitehead, T. A. Comprehensive sequence-flux mapping of a levoglucosan utilization pathway in E. coli. ACS synthetic biology 2015, 4, 1235–1243.

