## Supplementary Information for "Disentangling the contribution of each descriptive characteristic of every single mutation to its functional effects"

October 23, 2020

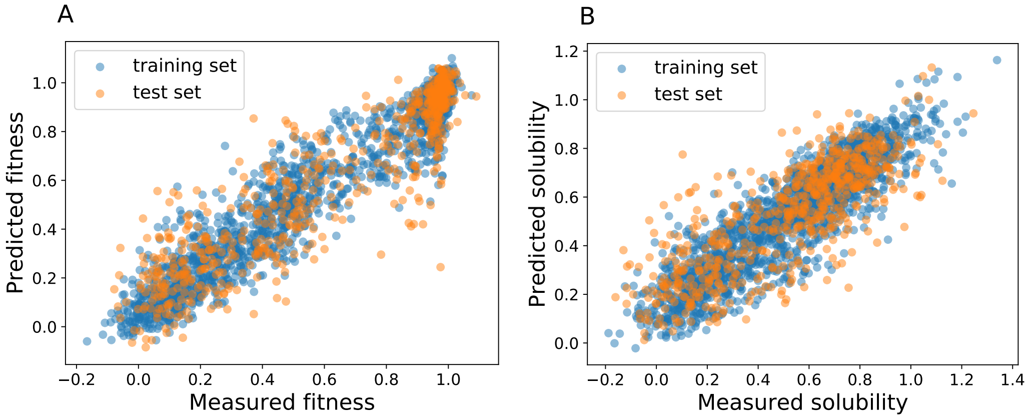


**Supplementary Figure 1**. **Quality of predictions.** A comparison of the predictions and observations from the deep mutational scan of β-lacatamase for (A.) fitness and (B.) solubility changes. The results obtained were in good agreement with the observations. 75% of the data was used for training.

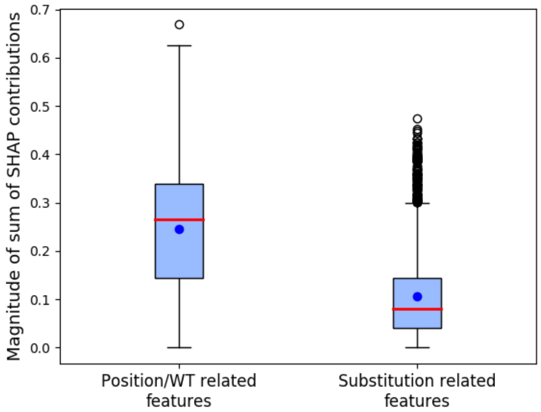


**Supplementary Figure 2**. **Relative contributions of groups of variables.** The relative contributions of the descriptive variables which mainly reflect the wild-type context are compared with the contributions from the descriptors which describe the precise nature of the mutation. The absolute values of the SHAP contributions from these two groups are shown. It is clear that more significant contributions arise from the wild-type context, rather than the precise mutations. An intuitive example is that a significant fraction of mutational effects can be understood from the knowledge of solvent accessibility or conservation of the amino acid being replaced.

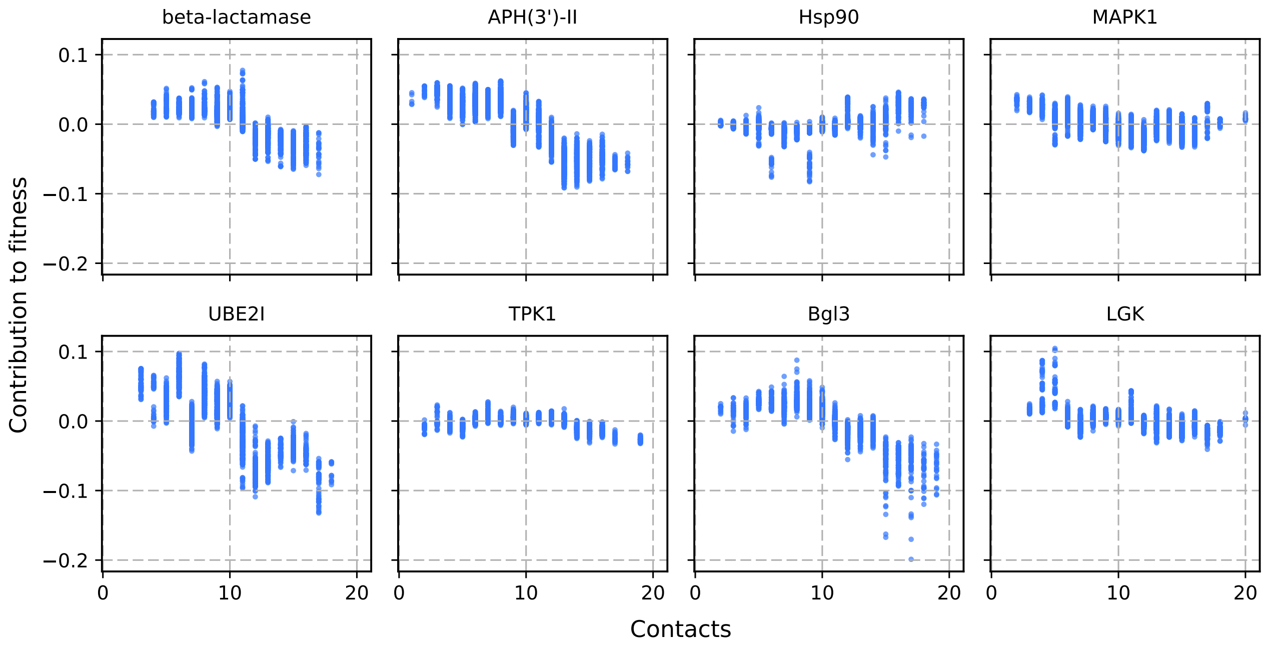


**Supplementary Figure 3. Role of number of wild-type contacts.** For the 8 proteins we analyzed, the contribution from the number of contacts a wild-type amino acid has with its neighboring amino acids, to the overall fitness changes observed in a mutational scan is shown.

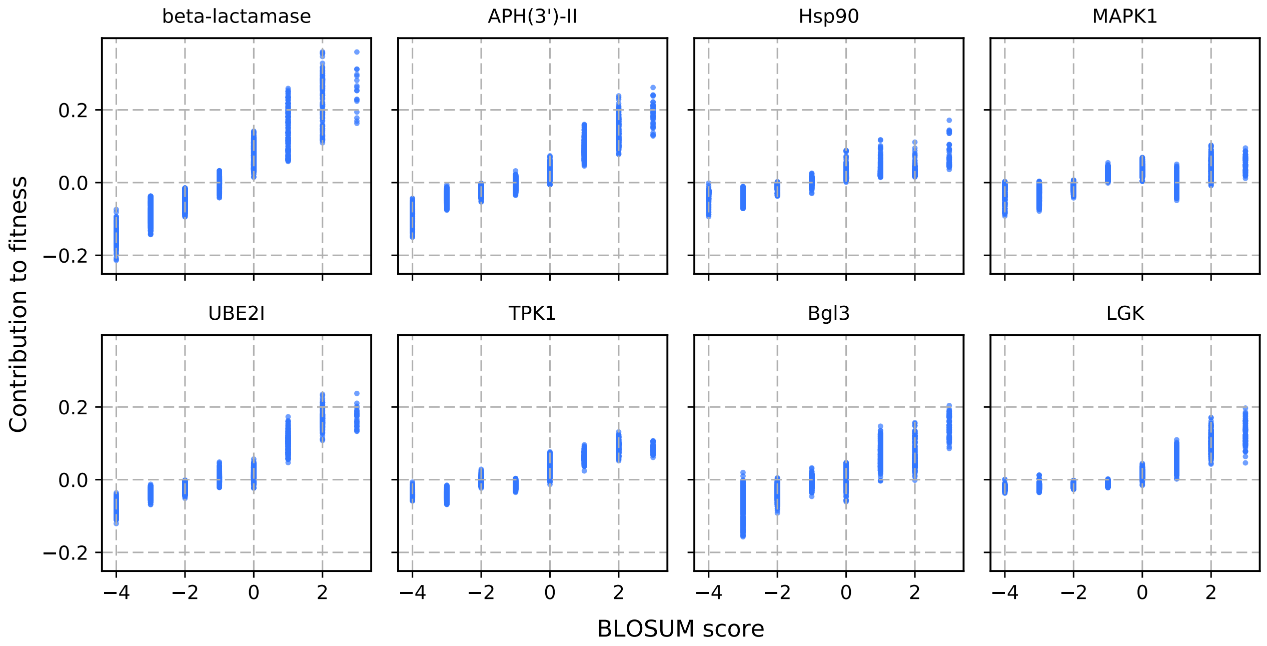


**Supplementary Figure 4. Role of substitution matrix score.** The substitution effects are generally described using BLOSUM substitution matrix.The analysis shows the degree of contribution from this descriptive factor to the overall mutational effects.

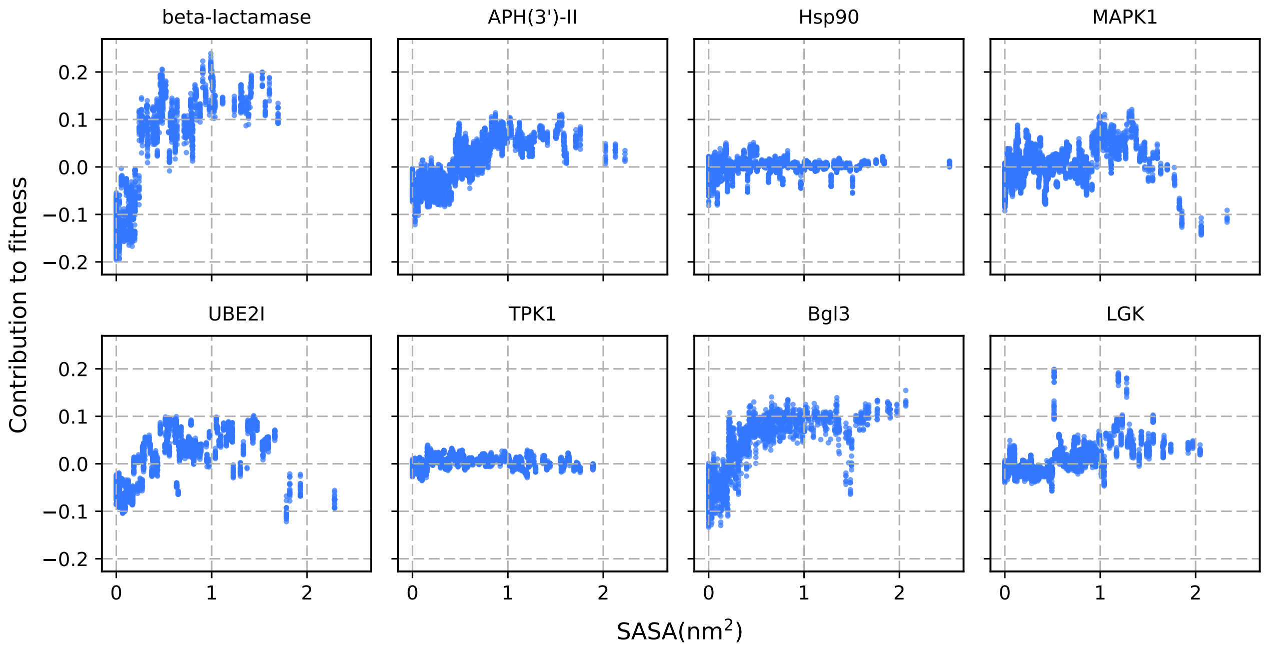


**Supplementary Figure 5. Role of solvent accessibility.** The contribution to the mutational effects from the solvent accessible surface area (SASA) is shown.

| Protein | No. of variants included in the analysis | Structure (PDB ID) | Reference |
|---|---|---|---|
| β-lactamase | 2467 | 1M40 | Stiffler et al.[1] |
| APH(3')-II | 4234 | 1ND4 | Melnikov et al.[2] |
| Hsp90 | 4021 | 2CG9 | Mishra et al.[3] |
| MAPK1 | 4470 | 4INF | Brenan et al.[4] |
| UBE2I | 2563 | 2UYZ | Weile et al.[5] |
| TPK1 | 3181 | 3S4Y | Weile et al.[5] |
| Bgl3 | 2732 | 1GNX | Romero et al.[6] |
| LGK | 6155 | 4ZLU | Klesmith et al.[7] |

**Supplementary Table** 1. Deep mutational scanning data sets used in the analysis.

| Step No. | Hyper parameters tuned | Parameter values sampled |
|---|---|---|
| 1 | *max_depth*, the maximum depth possible for each tree in the ensemble of trees | 3, 5, 7, 9, 15, 20, 25, 50 |
| | *min_child_weight*, the minimum number of samples that a node has in the tree | 1, 3, 5, 7, 9, 15, 20, 25, 50 |
| 2 | *gamma*, minimum loss reduction required for further division of a node of a tree | 0, 0.1, 0.2, 0.3, 0.4, 0.5, 0.6, 0.7, 0.8, 0.9 |
| 3 | *subsample*, fraction of total training samples that has to be used for growing a tree at each iteration | 0.4, 0.5, 0.6, 0.7, 0.8, 0.9 |
| | *colsample_bytree*, fraction of total features used at each iteration of training | 0.4, 0.5, 0.6, 0.7, 0.8, 0.9 |
| 4 | *reg_alpha*, weight for the L1 regularization term | 0, 0.0001, 0.001, 0.01, 0.1, 1, 10, 100 |

**Supplementary Table** 2. The steps in the hyper parameter optimization for the AI model. The values that were evaluated based on model predictive ability through a 5-fold cross-validation approach for each parameter are also given.

| Protein | n_estimators | max_depth | min_child_weight | gamma | subsample | colsample_bytree | alpha | RMSE$_{training}$ | RMSE$_{test}$ | $\rho_{training}$ | $\rho_{test}$ |
|---|---|---|---|---|---|---|---|---|---|---|---|
| **β-lactamase (solubility)** | 1116 | 5 | 3 | 0 | 0.8 | 0.7 | 0.0001 | 0.12 | 0.18 | 0.9 | 0.79 |
| **β-lactamase (fitness)** | 1247 | 9 | 25 | 0 | 0.8 | 0.8 | 0 | 0.1 | 0.15 | 0.96 | 0.91 |
| **APH(3')-II** | 1253 | 7 | 15 | 0 | 0.8 | 0.4 | 0.01 | 0.14 | 0.2 | 0.89 | 0.77 |
| **Hsp90** | 1218 | 5 | 1 | 0 | 0.6 | 0.6 | 0 | 0.1 | 0.14 | 0.93 | 0.86 |
| **MAPK1** | 1550 | 7 | 25 | 0 | 0.7 | 0.4 | 0.001 | 0.14 | 0.18 | 0.88 | 0.79 |
| **UBE2I** | 918 | 7 | 20 | 0 | 0.9 | 0.4 | 0.01 | 0.17 | 0.21 | 0.85 | 0.72 |
| **TPK1** | 276 | 7 | 7 | 0 | 0.7 | 0.4 | 0.001 | 0.23 | 0.27 | 0.68 | 0.34 |
| **Bgl3** | 660 | 7 | 5 | 0 | 0.9 | 0.7 | 0.001 | 0.11 | 0.18 | 0.92 | 0.75 |
| **LGK** | 804 | 7 | 7 | 0.1 | 0.7 | 0.4 | 0 | 0.18 | 0.2 | 0.69 | 0.52 |

**Supplementary Table** 3. The optimized set of hyper parameters for the different proteins are shown. Also shown are the quality of the results described by the root mean squared error (RMSE) and the Pearson-correlation ($\rho$) for the training and test sets.

| Protein | number of amino acids | number of domains | domain | % of residues in helices | % of residues in beta-sheet | % of charged and polar residues | % of hydrophilic surface area |
|---|---|---|---|---|---|---|---|
| β-lactamase | 286 | 1 | 1 | 46 | 17 | 45 | 77 |
| APH(3')-II | 264 | 1 | 1 | 39 | 18 | 43 | 65 |
| Hsp90 | 709 | 3 | ATPase binding domain | 32 | 16 | 55 | 72 |
| MAPK1 | 360 | 1 | 1 | 38 | 13 | 47 | 68 |
| UBE2I | 159 | 1 | 1 | 35 | 19 | 46 | 67 |
| TPK1 | 243 | 2 | both | 26 | 30 | 47 | 67 |
| Bgl3 | 500 | 1 | 1 | 43 | 16 | 43 | 73 |
| LGK | 447 | 1 | 1 | 42 | 21 | 46 | 72 |

**Supplementary Table** 4. Different protein descriptors that were analyzed to check for the possible correlation with variations in the individual variable contributions to fitness across proteins.

### References


[1] M. A. Stiffler, D. R. Hekstra, and R. Ranganathan, "Evolvability as a Function of Purifying Selection in TEM-1 beta-Lactamase," *Cell*, vol. 160, pp. 882–892, FEB 26 2015.

[2] A. Melnikov, P. Rogov, L. Wang, A. Gnirke, and T. S. Mikkelsen, "Comprehensive mutational scanning of a kinase in vivo reveals substrate-dependent fitness landscapes," *Nucleic Acids Research*, vol. 42, no. 14, 2014.

[3] P. Mishra, J. M. Flynn, T. N. Starr, and D. N. A. Bolon, "Systematic Mutant Analyses Elucidate General and Client-Specific Aspects of Hsp90 Function," *Cell Reports*, vol. 15, pp. 588–598, APR 19 2016.

[4] L. Brenan, A. Andreev, O. Cohen, S. Pantel, A. Kamburov, D. Cacchiarelli, N. S. Persky, C. Zhu, M. Bagul, E. M. Goetz, A. B. Burgin, L. A. Garraway, G. Getz, T. S. Mikkelsen, F. Piccioni, D. E. Root, and C. M. Johannessen, "Phenotypic Characterization of a Comprehensive Set of MAPK1/ERK2 Missense Mutants," *Cell Reports*, vol. 17, pp. 1171–1183, OCT 18 2016.

[5] J. Weile, S. Sun, A. G. Cote, J. Knapp, M. Verby, J. C. Mellor, Y. Wu, C. Pons, C. Wong, N. van Lieshout, F. Yang, M. Tasan, G. Tan, S. Yang, D. M. Fowler, R. Nussbaum, J. D. Bloom, M. Vidal,

D. E. Hill, P. Aloy, and F. P. Roth, "A framework for exhaustively mapping functional missense variants," *Molecular Systems Biology*, vol. 13, DEC 2017.

[6] P. A. Romero, T. M. Tran, and A. R. Abate, "Dissecting enzyme function with microfluidic-based deep mutational scanning," *Proceedings of the National Academy of Sciences*, vol. 112, no. 23, pp. 7159–7164, 2015.

[7] J. R. Klesmith, J.-P. Bacik, R. Michalczyk, and T. A. Whitehead, "Comprehensive sequence-flux mapping of a levoglucosan utilization pathway in e. coli," *ACS synthetic biology*, vol. 4, no. 11, pp. 1235–1243, 2015.
